# Systematic search for schizophrenia pathways sensitive to perturbation by immune activation

**DOI:** 10.1101/730655

**Authors:** Yue Gao, Xiaozhen Liang, Zhonglu Ren, Yanjun Li, Xinping Yang

**Affiliations:** Department of Obstetrics and Gynecology, Nanfang Hospital, Southern Medical University, Guangzhou 510515, China; Key Laboratory of Mental Health of the Ministry of Education, Southern Medical University, Guangzhou 510515, China; Department of Bioinformatics, School of Basic Medical Sciences, Southern Medical University, Guangzhou 510515, China

**Keywords:** Complex disease, Schizophrenia, disease candidate genes, immune genes, protein-protein interaction network, neighborhood walking, disease network, synapse remodeling

## Abstract

Immune activation has been recently found to play a large part in the development of schizophrenia, but the underlying mechanism remains largely unknown. Here, we report the construction of a high-quality protein interaction network for schizophrenia (SCZ Network) using a “neighborhood walking” approach to searching across human interactome network for disease-related neighborhoods. The spatiotemporal expression pattern of the immune genes in the SCZ Network demonstrates that this disease network is sensitive to the perturbation of immune activation during mid- to late fetal development and adolescence. The immune genes in the network are involved in pathways regulating the formation, structure and function of synapses and neural connections. Using single-cell transcriptome sequencing on the brains of immune-activated mice, we found that immune activation disturbed the SCZ network in the major brain cell types and the dysregulated pathways were also involved in synapse regulation, demonstrating that our “neighborhood walking” approach enables biological discovery in complex disorders.

## Introduction

Schizophrenia (SCZ) has a life time incidence rate around 1%, and is associated with disability of the patients and severe socioeconomic burden for families and society^1,2^. It is a polygenic complex disorder with a wide range of psychotic symptoms^3,4^. These include positive symptoms such as delusions, hallucinations and disorganized speech or behavior, negative symptoms such as loss of emotional expression or motivation, and cognitive symptoms such as deficits in memory, attention, working memory, processing speed and problem solving.

Schizophrenia has a heritability of about 80%^5,6^ and its genetic risk factors show very high heterogeneity. Functional genetics studies have identified several schizophrenia susceptibility genes such as DISC1^7^, PDE4B^8^, NPAS3^9^ and NRG1^10^. Many SCZ-associated loci have been identified through genome-wide association studies (GWASs)^11,12,13,14^. A recent large-scale GWAS by PGC (Psychiatric Genomics Consortium) has identified 108 loci significantly associated with schizophrenia^15^. The major histocompatibility complex (MHC)^16–19^ has been reproducibly found to be associated with schizophrenia. Some hotspots of genomic deletions or duplications have been identified at 1q21.1, 15q11.2 and 15q13.3, 22q11.2 and 16p11.2^20,21^. In addition, accumulating SCZ candidate genes have been found by whole exome sequencing^22,23^. Most notably, a large-scale study on exome sequencing of schizophrenia patients has discovered de novo mutations enriched in glutamatergic postsynaptic proteins comprising activity-regulated cytoskeleton-associated protein (ARC) and N-methyl-D-aspartate receptor (NMDAR) complexes^22^, suggesting dynamic synapse regulation may play a role in the disease development.

Accumulating evidence points to the important role of abnormal immune activation in the etiology of schizophrenia. Proinflammatory cytokine levels have been consistently reported to be increased in the serum of schizophrenia patients^24,25^ and also in the prefrontal cortex^26,27^. Prenatal exposure to maternal immune activation increases the risk of schizophrenia later in adult life^28–30^. A recent study found increased microglia-mediated synaptic pruning in the patient-derived neural culture^31^. Genome-wide association studies have identified loci covering genes involved in immune function^16–19^. Among the 3,437 SCZ candidate genes we collected from databases and literature, about 20% (714/3,437) of them are immune genes. Since the immune system is very sensitive to environmental insults, the fact that such a big portion of immune genes are found among the SCZ candidate genes suggests that immune activity may play a key role in the pathogenesis of schizophrenia.

Proteome-scale studies of protein interactions have shed light on molecular etiology of diseases and putative therapeutic targets^32,33^. High throughput mapping of protein interactions^34–36^ and literature curation of small-scale studies^37^ have accumulated huge amount of protein-protein interaction data. Recently, the interactions from different sources have been integrated into a comprehensive scored human interactome network containing about 600,000 interactions^38^. How to utilize the huge amount of protein interaction data to search for pathways underlying complex disorders such as schizophrenia remains challenging, because the candidate genes identified from genome-scale studies include a considerable portion of false positives. An efficient way to identify the true causal genes and the molecular pathways in which they are involved is in urgent need.

Here, we report our construction of a high-quality protein interaction a molecular network for schizophrenia (SCZ Network) by searching for disease-associated neighborhoods across a comprehensive human interactome (which we termed as “neighborhood-walking”). By integrating the spatiotemporal gene expression data of human brain, we have identified two developmental stages when the SCZ Network is sensitive to the perturbation of immune activation and the pathways that potentially link immune activation to schizophrenia through regulating the formation, structure and function of synapses. To verify that these potential schizophrenia molecular pathways that are sensitive to the perturbation of immune activation, we generated an immune-activated mouse model and carried out single-cell sequencing on the brain. By integrating the single-cell transcriptome sequencing data with the SCZ Network, we found that the immune activation-induced genes were also involved in pathways regulating synapse remodeling.

## Results

### Extracting a protein-protein interaction network for schizophrenia (SCZ Network) using the “neighborhood walking” method

We collected 3,437 SCZ candidate genes from databases and literature, and about 21% (714/3,437) of them are immune genes (Supplementary Data 1). Despite the large number of genes involved in SCZ, patients display similar core phenotypic features, suggesting that SCZ risk genes may share a common molecular network. Therefore, in order to remove the functionally unrelated false positives of disease candidate genes, identify causal genes and construct a molecular network for schizophrenia, we developed a “neighborhood walking” approach to searching for SCZ-related neighborhoods in a comprehensive human interactome network, which contains 17,210 genes and about 600,000 interactions^38^ (Fig. 1).

**Fig. 1.**
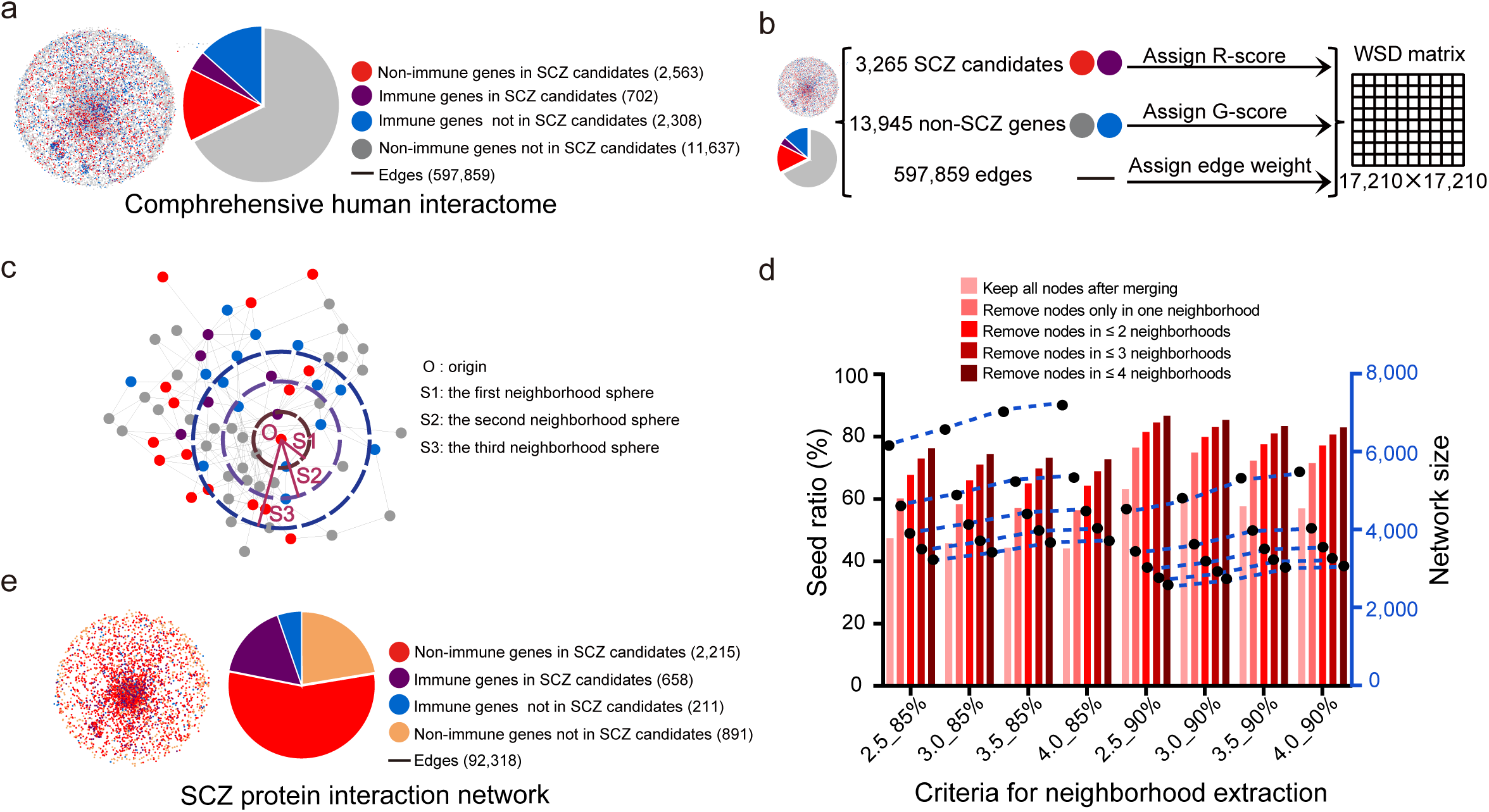
Extracting the protein interaction network for schizophrenia (SCZ Network) using “neighborhood walking” across the human interactome network. a. Mapping SCZ candidate genes (as seeds) to the comprehensive human interactome network containing 17,210 nodes (3,265 seeds; 13,945 nonseeds) and 597,859 edges. See also Supplementary Data 2.
b. Assignment of weight to nodes and edges: (1) For each seed, an R-score (risk score) was assigned based on how many times the seed appeared in literature or databases (R-score = 1-1.4^−x^, x is the count of times the gene appears in databases and literature); (2) For each nonseed, a G-score (guilt-by-association score) was assigned based on the average R-score of its interactors; (3) For each edge, the sum of the R-score and/or G-score of the two connecting nodes was taken as edge weight (W); (4) Weighted shortest distance (WSD) was calculated between any node pairs as 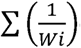 (Wi is the weight of an edge in a given shortest path). See also Supplementary Data 2.
c. “Neighborhood walking” method for extracting candidate networks for schizophrenia: (1) The “walking” was started from a seed (origin) and moved along the edges outward; (2) The neighborhood borders were defined by spheres with a WSD 0.5 in between; (3) The “walking” would stop if further “walking” would make the seed ratio of the neighborhood drop significantly (*p* < 0.05); (4) The neighborhoods with seed ratio cutoff at 85% or 90% were selected as SCZ-related neighborhoods, and thus were merged to obtain candidate networks for further consideration; (5) The nodes which appear 1, 2, 3 or 4 times or fewer (F≤1, 2, 3 or 4) in the selected neighborhoods were removed from the candidate networks.
d. Comparison of the seed ratios (fractions of seed nodes in the network) and network sizes of the candidate SCZ networks. The bars represent seed ratios and the blue curves represents the network sizes. The criteria for neighborhood extraction are shown as “WSD_x%” on the X-axis: the maximum neighborhood walking distances (WSD 2.5, 3.0, 3.5 or 4.0) and the seed ratios (85% or 90%). The colors of the bars represent the removal of nodes which appear less than 1, 2, 3 or 4 times (F≤1, 2, 3 or 4) in selected neighborhoods. See also Supplementary Data 3.
e. The SCZ protein interaction network (SCZ Network) and the nodes split as immune genes vs. non-immune genes or SCZ candidate genes vs. non-SCZ-candidate genes. See also Supplementary Data 4.

The “neighborhood walking” approach searches for disease-related neighborhoods in a human interactome network, based on the presumption that functionally related genes are more likely to cluster together in the molecular network. We hypothesized that, in a polygenic disease such as schizophrenia, the causal genes would interact with each other at molecular level to determine the complex traits. Based on this hypothesis, the causal genes would be more likely to remain in the selected SCZ-related neighborhoods, whereas the false positives of the disease candidate genes would tend to fall outside. We constructed a high-quality SCZ Network using “neighborhood walking” approach as following: (i) we first mapped the SCZ candidate genes as “seeds” onto the human interactome network (Fig. 1a, Supplementary Data 2); (ii) for each seed, we assigned a risk score (R-score = 1-1.4^−x^, x is the count of times the gene appears in databases and literature), and for each nonseed, we calculated a guilt score (G-score = the average R-score of its interactors) (Supplementary Data 2); (iii) for each edge, we calculated an edge weight (W = the sum of the R-score and/or G-score of the two interacting nodes) and a weighted edge length (WEL) 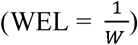 (Supplementary Data 2); (iv) for any two nodes in the network, we calculated a weighted shortest distance (WSD): 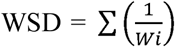 (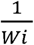 is the weighted edge length of an edge in a given shortest path) (Fig. 1b); (v) for each seed as an origin, we mapped the neighborhood spheres with WSD 0.5 in between to determine the borders of the neighborhood (Fig. 1c); (vi) we started the “walking” from an origin and stopped at a sphere if further “walking” to next sphere made the seed ratio in the neighborhood drop significantly; (vii) we used cutoffs of seed ratio at 85% or 90% and cutoffs of maximum walking distance at WSD 2.5, 3.0, 3.5 and 4.0 for selecting SCZ-related neighborhoods (Supplementary Data 3); (viii) we then merged the SCZ-related neighborhoods to obtain candidate SCZ networks (Fig. 1d); (ix) For each candidate SCZ network, we removed the nodes only appears fewest times (cutoffs for frequency: F ≤ 1, 2, 3 or 4) in selected neighborhoods and then compared the candidate SCZ networks using seed ratios as indicatives of relevance to the disease (Fig. 1d, Supplementary Data 3).

The seed ratio cutoff at 90% for selecting neighborhoods gave better performance than the seed ratio cutoff at 85%, while the different cutoffs of maximum walking distance at WSD 2.5, 3.0, 3.5 and 4.0 gave similar outcomes based on seed final ratios of the networks (Fig.1d). The increased cutoffs of maximum walking distance expanded network sizes, and the size expansion reached a stationary phase at cutoff of maximum walking distance WSD 3.5 (Fig. 1d). The removal of the nodes appearing only once (F ≤ 1) in the selected neighborhoods gave the biggest improvement of final seed ratios in the networks (Fig. 1d). Based on these results, we chose the following criteria for SCZ network construction: the cutoff of seed ratio at 90% and the cutoff of maximum walking distance at WSD 3.5 for selecting SCZ-related neighborhoods, and the cutoff of the node frequency at F ≤ 1 in the selected neighborhoods for removing nodes from the candidate SCZ networks. Using on these optimal cutoffs, we obtained a SCZ Network, which has a seed ratio of 72.3% and contains 3,975 nodes and 92,318 edges, including 869 immune genes (658 known SCZ candidate genes) and 3,106 non-immune genes (2,215 known SCZ candidate genes) (Fig. 1e, Supplementary Data 4).

### The high quality SCZ network

The primary purpose of our “neighborhood walking” approach is to extract a high-quality disease network from human interactome by removing false positives of SCZ candidate genes while keeping the true positives and novel candidate genes in the network. Of the 3,437 SCZ candidate genes, 3,265 were mapped on the human interactome (Fig. 2a). After the “neighborhood walking” process, 2,873 schizophrenia candidate genes and 1,102 novel candidate genes were kept in the SCZ Network (Fig. 1e). In order to verify that the “neighborhood walking” approach has removed false positives and kept true positives, we evaluated the SCZ relevance of the kept genes (genes inside the SCZ Network) and removed genes (genes outside the SCZ Network) using three datasets collected from literature (Supplementary Data 5): (i) dysregulated genes in the brain of schizophrenia patients^39–42^, (ii) variants associated with schizophrenia^43^ and (iii) genes with brain-specific expression (Expression Atlas)^44^. Compared to removed genes, the kept genes showed significant enrichment with dysregulated genes in the brains of schizophrenia patients (*p* = 0.0007 for all kept genes, two-side Fisher’s exact tests) (Fig. 2b, Supplementary Data 5), SCZ-associated variants (*p* = 5.9827e-98 for all kept genes, *p* = 7.5082e-07 for kept seeds, *p* = 0.0183 for kept nonseeds, two-side Fisher’s exact tests) (Fig. 2c, Supplementary Data 5), and brain-specific genes (*p* = 2.3283e-99 for all kept genes, *p* = 6.0824e-11 for kept nonseeds, two-side Fisher’s exact tests) (Fig. 2d, Supplementary Data 5). Removed seeds showed depletion of dysregulated genes (Fig. 2b), SCZ-associated variants (Fig. 2c) and brain specific genes (Fig. 2d), suggesting that some false positives were removed. The kept nonseeds showed significant enrichment with SCZ-associated variants (Fig. 2c) and brain-specific genes (Fig. 2d), suggesting that they could be novel SCZ candidate genes. Among the 16 tissues, the kept genes showed increased brain-specific expression compared with the genes in the whole human interactome network and the genes removed (ratio = 10.6% for all genes in the human interactome, 20.3% for all kept genes in the SCZ Network, 7.7% for removed genes), but not in other tissues (Fig. 2e, Supplementary Data 5).

**Fig. 2.**
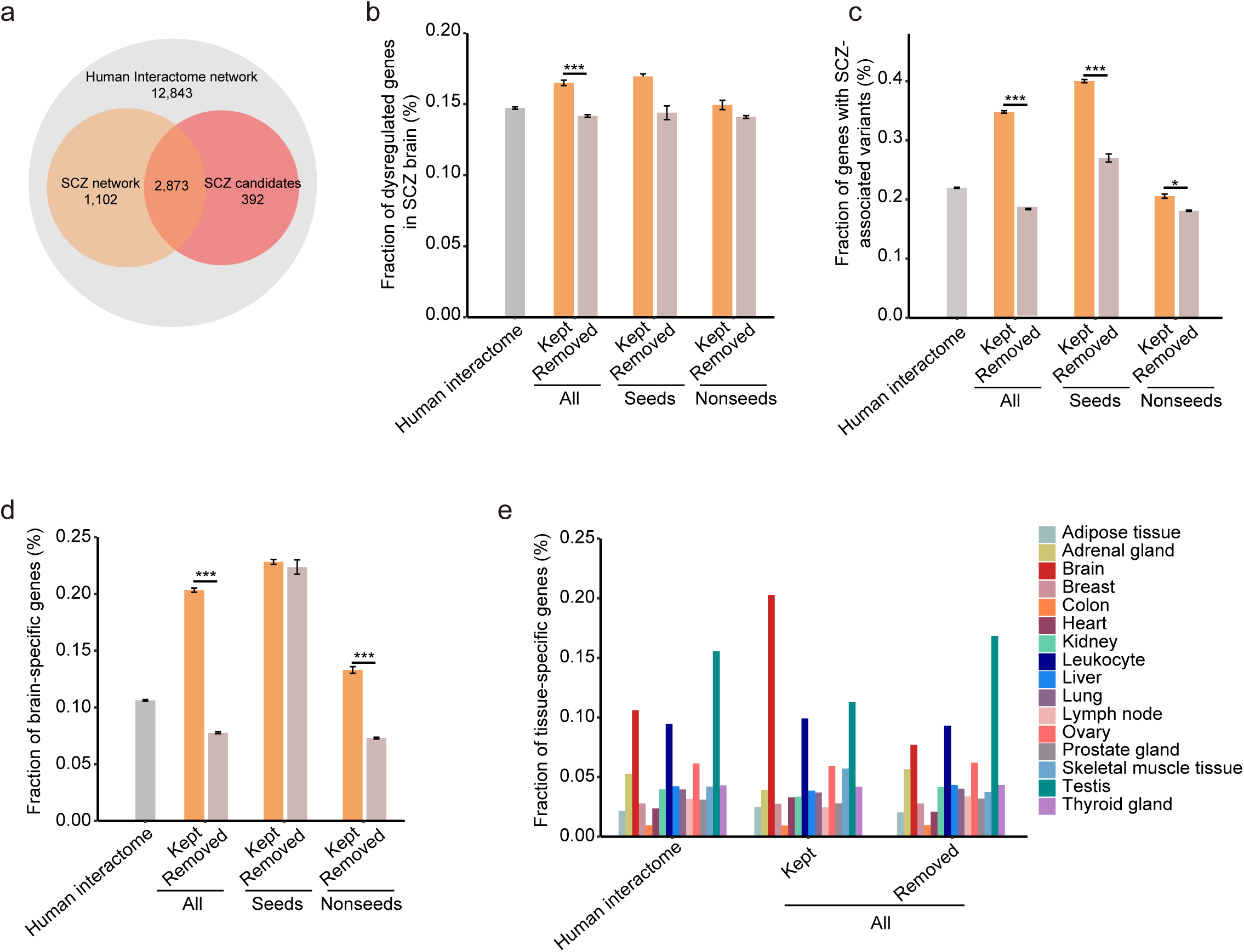
The high quality SCZ Network. a. Venn diagram of the genes in the SCZ Network (orange), SCZ candidate genes as seeds (red) and the genes in the human interactome (grey).
b. Fraction of SCZ-brain dysregulated genes in the SCZ Network (kept genes), outside the SCZ Network (removed genes) and in the human interactome.
c. Fraction of genes with SCZ-associated variants in the SCZ Network (kept genes), outside the SCZ Network (removed genes) and in the human interactome.
d. Fraction of brain-specific genes in the SCZ Network (kept genes), outside the SCZ Network (removed genes) and in the human interactome.
e. Tissue-specificity of genes in the SCZ Network (kept genes), not in the SCZ Network (removed genes) and in human interactome. Y-axis: fraction of tissue specific genes in the network.

Two-sided Fisher’s exact test was used to compare the kept nodes with the removed nodes, \**p* < 0.05, \*\*\**p* < 0.001. Error bars represent the standard error of the fraction, estimated using bootstrapping method with 100 resamplings.

### Two sensitive stages for SCZ Network to be disturbed by immune activation

The SCZ Network contains 869 (21.9%) immune genes and 3,106 non-immune genes (Fig. 1e). Consistent with recent studies which discovered the involvement of immune system in the development of schizophrenia^50,51,26,27–29,28–30^, we found significant enrichment of immune genes in the SCZ Network (*p* = 5.1042e-16, two-side Fisher’s exact test) (Fig. 3a). We wondered if the spatiotemporal dynamics of the SCZ Network might reveal the sensitive stages and brain regions to immune activation, and more importantly, the underlying molecular pathways. We utilized the expression data from 4 regions of human brains at 15 different developmental periods^45,46^ to calculate the overrepresentation of the genes in the SCZ Network in different spatiotemporal phases (Fig. 3b) (see Methods for determining phase-specific genes). The *p*-values for the enrichments were calculated using one-side Fisher’s exact test. Region 1 includes: posterior inferior parietal cortex, primary auditory (A1) cortex, primary visual (V1) cortex, superior temporal cortex, inferior temporal cortex; region 2 includes: primary somatosensory cortex, primary motor cortex, orbital prefrontal cortex, dorsolateral prefrontal cortex, medial prefrontal cortex and ventrolateral prefrontal cortex; region 3 includes: striatum, hippocampus and amygdala; region 4 includes: mediodorsal nucleus of the thalamus and cerebella cortex (Fig. 3b).

**Fig. 3.**
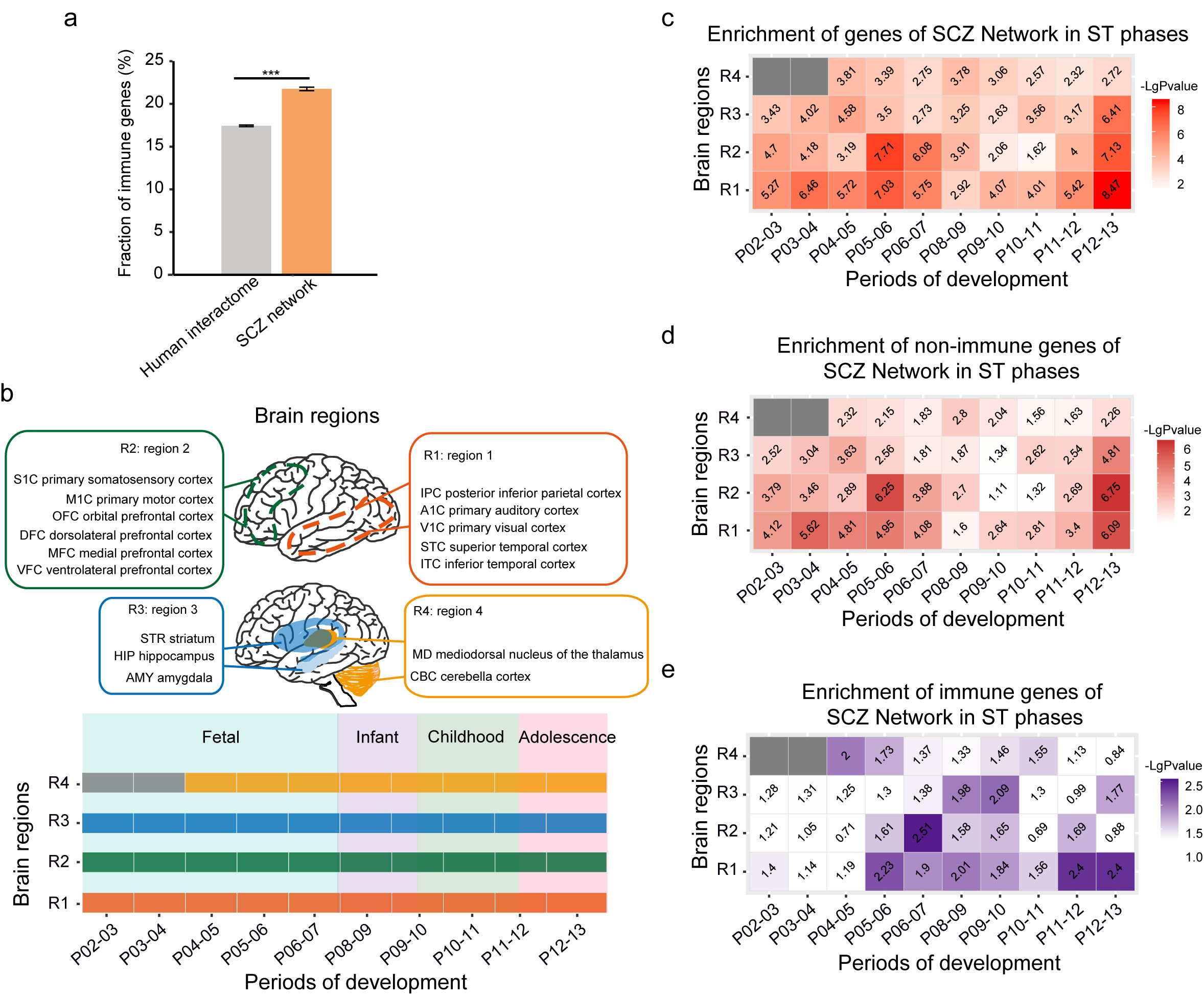
Spatiotemporal dynamic expression of the genes in the SCZ Network during brain development. a. Enrichment of immune genes in the SCZ Network (*p* = 5.1042e-16, two-side Fisher’s exact test). Error bars represent the standard error of the fraction, estimated using bootstrapping method with 100 resamplings, \*\*\**p* < 0.001.
b. A schematic of human spatiotemporal expression data source: 4 brain regions and 8 developmental periods (see also Methods for a list of the periods).
c. Enrichment of the genes of the SCZ Network in spatiotemporal phases. The heatmap shows the significance levels of overrepresentation of the genes of the SCZ Network in the spatiotemporal phases.
d. Enrichment of the non-immune genes of the SCZ Network in spatiotemporal phases. The heatmap shows the significance levels of overrepresentation of the non-immune genes of the SCZ Network in the spatiotemporal phases.
e. Enrichment of the immune genes of the SCZ Network in spatiotemporal phases. The heatmap shows the significance levels of overrepresentation of the immune genes of the SCZ Network in the spatiotemporal phases.

One-sided Fisher’s exact test was used for the analyses of enrichments. The numbers in the squares represent the -log (*p-*value). “ST” stand for “spatiotemporal” in panel c, d and e. See also Supplementary Data 6 for detailed information.

The genes in the SCZ Network are more enriched in region 1 during periods 2-7, 11-13 (early- to late fetal development, and adolescence to young adulthood, respectively), region 2 during periods 5-7 (early-mid fetal to late fetal development) and 12-13 (adolescence to young adulthood) and region 3 during periods 12-13 (Fig. 3c). Non-immune genes in the SCZ Network show a similar pattern as all genes in the network (Fig. 3d).

To further identify the brain regions and developmental stages that are sensitive to immune activation, we further calculated the overrepresentation of the immune genes of SCZ Network in the spatiotemporal phases. The immune genes are more enriched in region 1 during periods 5-6 (mid fetal development) and 11-13 (childhood to young adulthood); in region 2 during periods 6-7 (late-mid to late fetal development); in region 3 during periods 9-10 (late infancy to early childhood) (Fig. 3e). The immune system has evolved to response to environmental changes and protects organisms from infection and injury through immune responses. Immune genes in the SCZ Network show different spatiotemporal expression pattern from non-immune genes during fetal development (Fig. 3d-3e). Non-immune genes in the SCZ Network are more likely to be expressed in region 1 and 2 (cortex regions) during both early fetal and mid fetal development, while immune genes are more likely to be expressed from mid fetal to late fetal development. Both non-immune genes and immune genes in the SCZ Network are more likely to be expressed in region 1 at adolescence to early adulthood.

The overrepresentation of the genes in the SCZ Network in the stages during both early- to late development and adolescence suggests that this network may be more susceptible to perturbation during the whole fetal development and adolescence (Fig. 3c). The overrepresentation of immune genes in this network in the stages during both mid- to late fetal development and adolescence (Fig. 3e) suggests that it may be more easily affected by immune activation during both mid- to late fetal development and adolescence.

### The immune genes are involved in molecular network linking immune activation to synapse remodeling

In order to further investigate how the immune genes might be involved in the molecular mechanism underlying schizophrenia, we extracted an immune-centered subnetwork (SCZ-i Subnetwork) (Fig. 4a), containing 869 immune genes and their 2,924 first-degree neighbors, from the SCZ Network. The 2, 924 non-immune genes interact with the immune genes suggest that their functions may be regulated by the immune genes. These genes in the subnetwork tend to be more expressed in early fetal developmental stage (P2-4), mid-late fetal developmental stage (P5-6) and adolescence to young adult stage (P12-13) (Fig. 4b), suggesting that these genes are more active during these three developmental stages.

**Fig. 4.**
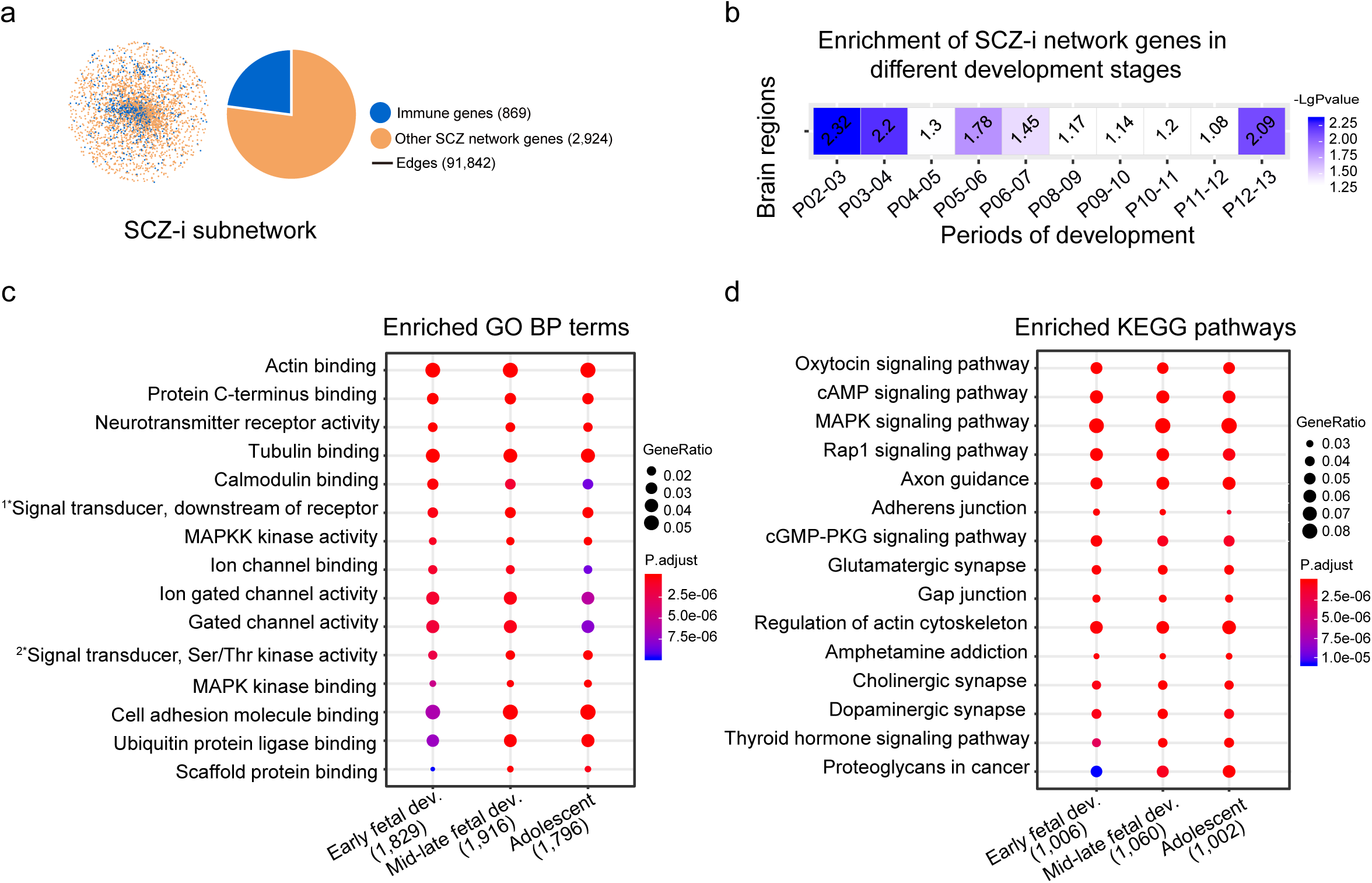
SCZ-immune Subnetworks (SCZ-i Subnetworks) of the brain at early fetal stage, mid fetal stage and adolescent stage. a. Construction of the SCZ-immune Subnetwork (SCZ-i Subnetwork): immune genes were mapped onto the SCZ Network and the SCZ-i Subnetwork was extracted to include the immune genes and their first-degree neighbors.
b. Enrichment of the genes of the SCZ-i Subnetwork in the brain (merged from 4 regions) across 8 developmental periods.
c-d. Functional enrichment analyses on the SCZ-i Subnetworks for early fetal developmental stage, mid-late fetal developmental stage and adolescent stage (See also Supplementary Fig. 1 for the SCZ-i subnetworks at P02-03, P05-06 and P12-13). Top 15 highly enriched GO BP terms (c) or KEGG pathways (d) are shown as dots. The enriched GO terms or KEGG pathways are those with overrepresentation of genes in the SCZ-i Subnetworks. The sizes of the dots represent gene ratios and the colors represent the adjusted *p*-values. The numbers under each stage are the numbers of identified genes in each category. The full terms for the two abridged GO terms: 1* signal transducer activity, downstream of receptor; 2* signal transducer, downstream of receptor, with serine/threonine kinase activity. See also Supplementary Data 7 for complete information of enriched GO terms and KEGG pathways.

To find out the neuronal functions these immune genes involved in, we extracted SCZ-i Subnetworks for these three developmental stages by mapping the stage co-expressed genes on the SCZ-i Subnetwork (Supplementary Fig. 1) and further performed functional enrichment analyses to search for functions of these subnetworks (Fig. 4c-d). The enriched GO terms include activity of cytoskeleton components (actin binding, tubulin binding and scaffold protein binding), ion channel binding/activity, neurotransmitter activity, MAPKK binding and MAPKKK activities (Fig. 4c, Supplementary Data 7). The enriched pathways include regulation of actin cytoskeleton, adhesion junction, gap junction, axon guidance, glutamatergic synapse, dopaminergic synapse, oxytocin signaling, MAPK signaling, estrogen signaling and cAMP signaling pathways (Fig. 4d, Supplementary Data 7). Some of these pathways are obvious synapse signaling pathways such as axon guidance, glutamatergic synapse and dopaminergic synapse pathways, while some other pathways are reported to be involved in synapse regulation. The regulation of actin cytoskeleton is very important for remodeling and maintaining the structure and function of the synapses, because the spines are formed and supported by the complex dynamic cytoskeletal architectures of actin filaments^49^. Mutations that disrupt the actin filament structures can cause neurodevelopmental and psychiatric disorders such as schizophrenia^22,50^. The adhesion junction is important for synapse structure and stability^51,52^, and the gap junction is necessary for mediating electrical transmission^53^. Oxytocin signaling pathway has been known to regulate synapse structure, function and neuron connectivity during social interaction^54,55^. Recent studies have found that oxytocin signaling is involved in the differentiation of neuronal progenitor cells^56^, synapse formation^57^ and the formation of neural circuits during brain development^58,59^. MAPK signaling pathway^60,61^, estrogen signaling pathway^62,63^ and cAMP signaling pathway^64,65^ are important for synaptogenesis and synapse pruning and refining. Therefore, these results suggest that immune activation in the brain during early-, mid- to late fetal development and adolescence to young adulthood is more likely to interfere with pathways related to synapse formation, structure and remodeling. Consistent with this speculation, accumulating evidence demonstrates that prenatal maternal infections during pregnancy^28,29^ and infection events during adolescence are associated with schizophrenia^47,48^.

### Immune activation perturbs SCZ-related molecular pathways for synapse remodeling

In order to search for SCZ-related molecular pathways in different brain cell types affected by immune activation, we first activated the peripheral immune system by intraperitoneal injection (IPI) of lipopolysaccharide (LPS) into adolescent mice twice on post-natal day 45 and 46, and collected the serum from the mice on the third day. Then we performed intracerebroventricular injection (ICVI) of 15 ul of serum from the immune-activated mice (Fig. 5a, See Methods for details). After ICVI of immune-activated serum for one week, we carried out single-cell transcriptome sequencing on the brains from the mice. We used the *t*-stochastic neighbor embedding (*t*-SNE) method on high variable genes (HVGs)^66^ for separating cells into 9 clusters (Fig. 5b), and used markers to identify the cell types^67^ (Methods). We defined a gene as “expressed” if 5% of the cells had at least 1 read, and obtained expressed genes for each cell type (Supplementary Fig. 3, Supplementary Data 9). We defined differentially expressed genes (DEGs) for different cell types using R package monocle^68^. We found that DEGs in 4 major cell types (neuron, microglia, astrocyte and oligodendrocyte) were significantly enriched with genes in the SCZ Network, suggesting that these 4 cell types may be involved in the development of schizophrenia (Fig. 5c). After immune activation, these 4 cell types showed different extents of global gene expression changes, with 5,032 genes differentially expressed in neurons, 1,812 in microglia, 723 in astrocytes and 1,771 in oligodendrocytes (Fig. 5d, Supplementary Data 9). Most of the DEGs were different between these four cell types and only 137 DEGs were in common. We extracted subnetworks for DEGs in these 4 cell types from the SCZ Network, i.e. SCZ-micr-DEG Subnetwork, SCZ-astr-DEG Subnetwork, SCZ-olig-DEG Subnetwork and SCZ-neur-DEG Subnetwork, each containing cell-type-specific DEGs and their first neighbors in the SCZ network (Fig. 5e, Supplementary Fig. 3). Although DEGs in the 4 cell types are mostly different from each other (Fig. 5d), the DEG Subnetworks for the 4 cell types (Fig. 5e) are enriched in similar GO terms and KEGG pathways (Fig. 5f-g, Supplementary Data 9). Many of these enriched pathways, such as pathways in regulation of actin cytoskeleton, adherens junction, gap junction, axon guidance, glutamatergic synapse and dopaminergic synapse etc., were also found to be enriched with genes in SCZ-i Subnetwork as described above (Fig. 4d). These results confirm our findings on the SCZ Network that the perturbation of this network by immune activation may lead to synapse remodeling.

**Fig. 5.**
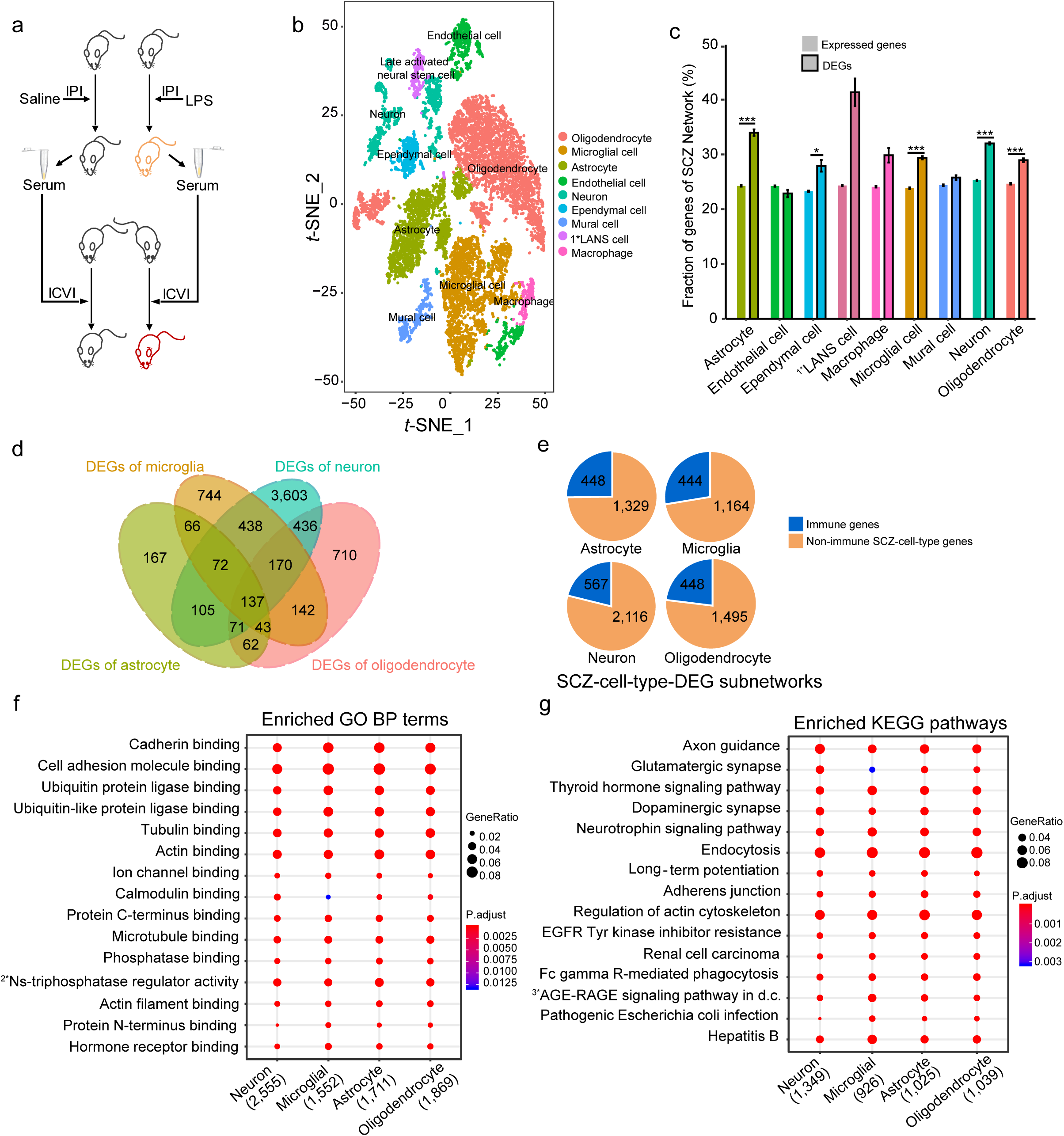
Singe-cell transcriptome profiling on the immune-activated brains of adolescent mice reveals key pathways linking immune activation to synapse remodeling. a. A schematic for the induction of immune activation in the brain (see also Methods).
b. The *t*-SNE plot constructed from the normalized and corrected logarithm of expression values of highly variable genes for cells. Cell-type clustering and identification: 11 subpopulations were identified and marked with different colors. (See Supplementary Fig. 3a for cell population clusters of the brain cells of mice with different treatments).
c. The enrichments of the genes of the SCZ Network in 9 different cell populations. Two-sided Fisher’s exact test was used for the enrichment analyses, \**p* < 0.05, \*\*\**p* < 0.001; error bars represent the standard error of the fraction, estimated using bootstrapping method with 100 resamplings (Supplementary Data 8).
d. Venn diagram of differentially expressed genes (DEGs) in 4 major cell types in the brain (Supplementary Data 8).
e. Genes in SCZ-DEG subnetworks for 4 major cell types (Neuron, microglia, astrocyte and oligodendrocyte split as immune genes (blue) and non-immune genes (red). These cell type-specific subnetworks of differentially expressed genes were extracted from the SCZ Network to include DEGs of the 4 cell types and their expressed first-degree neighbors (Supplementary Data 9).
f-g. The GO biological processes (BP) terms and KEGG pathways with overrepresentation of genes in 4 SCZ-DEG subnetworks for neuron, microglia, astrocyte and oligodendrocyte. Top 15 highly enriched GO BP terms (f) or KEGG pathways (g) are shown as dots. The sizes of the dots represent gene ratios and the colors represent the adjusted *p*-values. The numbers under each stage are the numbers of identified genes in each category. The full terms for the three abridged GO terms: 1* Late activated neural stem cell; 2* nucleoside-triphosphatase regulator activity; 3* AGE-RAGE signaling pathway in diabetic complications. See also Supplementary Data 9 for complete information of enriched GO terms and KEGG pathways.

## Discussion

The two most significant advances in searching for risk factors of schizophrenia over the last decade are: 1) large number of schizophrenia-associated genes have been identified by genome wide studies^15,22,23^; 2) immune activation has been repeatedly proven to be associated with this disease^24,25,26,27,28,29^. However, further search for underlying molecular mechanisms remains a big challenge. The “Two-Hit Hypothesis” of schizophrenia, proposed in 1999, suggest that schizophrenia needs two hits for its development: the first hit (basic requirement) includes genetic factors such as germline mutations and the second, environmental factors such as viral infection, pregnancy complications and social stressors^73,74^. Recently, it has been suggested that maternal immune activation may also serve as “the first hit”, which make the offspring more susceptible to a “second hit” during adolescence^30^. Immune activation has been reported to be much more involved in schizophrenia than we previously thought. Genome-wide association studies have repeatedly associated MHC locus with schizophrenia^15,16^. Among the 3,437 SCZ candidate genes we collected from databases and literature, about 714 (21%) of them are immune genes. Recently, the gene complement C4 is reported to be critical for synapse pruning^69^. For most of the immune genes, we still do not know how they link to schizophrenia at molecular level.

The large number of SCZ candidate genes are mostly from large-scale genomics studies, comprising considerable portion of false positives and false negatives. We have developed a “neighborhood walking” approach to identifying disease-related neighborhoods in a high-quality comprehensive human interactome network^38^. Using this approach, we have removed some false positives of the SCZ candidate genes which fall outside of the selected neighborhoods, and identified some novel candidate genes which fall within. By merging the SCZ-related neighborhoods, we were able to assemble a high-quality protein interaction network for schizophrenia (SCZ Network). Compared to the genes that fall outside of the SCZ Network (removed genes), the genes in the network (kept genes) are significantly enriched with SCZ-associated variants, dysregulated genes in the postmortem brains of SCZ patients and genes with brain-specific expression (Fig. 2). The removed SCZ candidate genes show significant depletion of SCZ-associated variants, while kept non-candidate genes show significant enrichment with SCZ-associated variants (Fig. 2). These results suggest that “neighborhood walking” may provide an effective approach for removing false positives of the disease candidate genes and identifying novel candidate genes.

In the SCZ Network, the immune genes interconnect with the non-immune genes (Fig. 1e). The spatiotemporal expression pattern of the genes in the SCZ Network (Fig. 3e) demonstrates that this molecular network is more active during both early- to late fetal development and adolescence and thus more likely to be disturbed during these developmental stages (Fig. 3c), and that the immune genes in the SCZ Network are more active during both mid- to late fetal development and adolescence (Fig. 3e). Since proinflammatory cytokine levels have been consistently reported to be increased in the serum of SCZ patients^24,25^ and also in prefrontal cortex^26^, immune activation are considered an important environmental risk factor. By extracting SCZ-i Subnetwork containing immune genes and their direct interactors in the SCZ Network, we have identified pathways in which the immune genes are involved, such as the pathways in regulation of actin cytoskeleton, adhesion junction, gap junction, axon guidance, glutamatergic synapse, dopaminergic synapse, and other signaling pathways that regulate synapse formation, structure and function (Fig. 4d). The enrichment of SCZ-i Subnetwork genes in these pathways suggest a pervasive participation of immune genes in synapse regulation during early fetal development, mid- to late fetal development and adolescence (Fig. 4c, 4d). These results are in consistent with previous findings that prenatal maternal infection greatly increase the risk for schizophrenia^28,29^, and adolescent stress may trigger the onset of this disease^70–72^.

Using single-cell transcriptome sequencing on the brain of mouse model of immune activation (Fig. 5a-b), we identified differentially expressed genes (DEGs) in each brain cell types. The overrepresentation of DEGs in microglia, astrocytes, oligodendrocytes and neurons in SCZ Network (Fig. 5c-d) suggests that these 4 cell types may be involved in the development of schizophrenia. It is well known that microglia and astrocytes carry out most of immune functions in the brain^31,75^. In the subnetworks of DEGs in the 4 major cell types extracted from SCZ Network (Fig. 5e, Supplementary Fig. 3), neurons and oligodendrocytes have similar or larger number of immune genes than those of astrocytes and microglia (Fig. 5e). This is surprising because it indicates that neurons and oligodendrocytes may also actively participate in the immune responses. We have identified similar pathways that are enriched with subnetworks of DEGs in the 4 major cell types. To our expectation, these pathways are mainly involved in synapse regulation, such as regulation of actin cytoskeleton, adherens junction, glutamatergic synapse and dopaminergic synapse (Fig. 5f-g), suggesting that these 4 cell types may participate in the immune activation-induced synapse remodeling.

We have shown that the disease network extracted from a comprehensive human interactome network using “neighborhood walking” approach has high quality. Although schizophrenia has thousands of candidate genes and there are considerable portions of false positives and false negatives, we have identified disease-related neighborhoods using “neighborhood walking” and assembled them into the SCZ Network. The spatiotemporal expression patterns of the genes in the SCZ Network have revealed sensitive developmental stages and brain regions, which are consistent with the clinical findings about the association of immune activation with schizophrenia and provide a molecular explanation for the “Two-Hit-Hypothesis” on the etiology. The identification of the immune gene-involved pathways suggests a pervasive role of immune activation in synapse regulation. Our analyses on single-cell transcriptome sequencing data of the brain from the mouse model, have confirmed that perturbation of the SCZ Network by immune activation leads to the dysregulated pathways in synapse remodeling.

## Acknowledgements

The work was supported by the Key Scientific and Technological Projects of Guangdong Province (2018B030335001), Science and Technology Projects of Guangzhou (Grant No. 201704020116), the National Natural Science Foundation of China (Grant No. 81571097), and the Natural Science Foundation of Guangdong Province (Grant No. 2016A030308020). The funding organizations had no role in design and conduct of the study; collection, management, analysis, and interpretation of the data; and preparation, review, or approval of the manuscript.

## Author contributions

X.Y. conceived the project; Y.G. performed the computational analysis; Z.R. participated in the analysis; Y.G., Y.L., X.L. acquired the data. All of the authors contributed to the preparation of the manuscript.

## METHODS

### 1. Collection of Schizophrenia candidate genes

A total number of 3,437 schizophrenia candidate genes (Supplemental Data 1) were collected from:

1. HGMD (The Human Gene Mutation Database, http://www.hgmd.cf.ac.uk/ac/index.php)^76^
2. SzGene (SchizophreniaGene, http://www.szgene.org/)^77^
3. SZDB (A database for Schizophrenia Genetic Research, http://www.szdb.org)^78^
4. SZGR (Schizophrenia gene resource, http://bioinfo.mc.vanderbilt.edu/SZGR/)^79^
5. PheGenI(Phenotype–Genotype Integrator, https://www.ncbi.nlm.nih.gov/gap/phegeni)^80^
6. SNPedia (https://www.snpedia.com/index.php/Schizophrenia)
7. Literature^81^

### 2. Collection of immune genes

A total number of 3,904 immune genes (Supplemental Data 1) were collected from:

1. InnateDB^82^ (Innate Immunity Genes, https://www.innatedb.com)
2. GO-immune-response (http://amigo.geneontology.org/amigo/term/GO:0006955);
3. ImmPort: (http://www.immport.org/immport-open/public/reference/genelists);
4. Nanostring’s Immune Profiling Panel Probe (https://www.nanostring.com/products/gene-expression-panels/gene-expression-panels-overview/hallmarks-cancer-gene-expression-panel-collection/pancancer-immune-profiling-panel?jumpto=SUPPORT);
5. IMGT/GENE-DB (the international ImMunoGeneTics information system): (http://www.imgt.org/genedb/).

### 3. Human interactome data

The comprehensive human interactome data were downloaded from InBioMap (https://www.intomics.com/inbio/map.html)^83^.

### 4. Weighted shortest distance

The SCZ candidate genes were mapped as “seeds” onto the human interactome network, shown as colored nodes in Fig. 1b. For each seed, a risk score (R-score) was assigned (R-score = 1-1.4^−x^, x is the count of times the gene appears in databases and literature); for each nonseed, a guilt score (G-score) was assigned (G-score = the average R-score of its interactors). For each edge, an edge weight was assigned (W = the sum of the R-score and/or G-score of the two interacting nodes). For any two nodes in the network, a weighted shortest distance (WSD) was calculated: WSD = 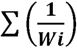 (W_i_ is weight of the edge in a given shortest path). Weighted shortest distance was calculated by R toolkit Igraph’s (version 1.2.4) distances function, using undirected graph and Dijkstra’s algorithm.

### 5. “Neighborhood walking” approach

For a given node as an origin, neighborhood spheres were mapped at WSD 2.5, 3.0, 3.5 and 4.0 as neighborhood borders, and the neighborhood was defined from an origin to a sphere. The “walking” started from an origin moving outwards on all directions and stopped at a sphere if further “walking” to next sphere made the seed ratio drop significantly. For each origin, only the neighborhood with biggest WSD (from WDS 2.5 to 4.0) was selected for further analyses. In order to select SCZ-related neighborhoods, the cutoffs for seed ratio were set as 85% or 90%, and the cutoffs for maximum walking distance were WSD 2.5, 30, 3.5 or 4.0 from origin to spheres. Since every seed was set as an origin, the selected SCZ-related neighborhoods should overlap with each other, and thus a node may appear in multiple neighborhoods. In order to remove false positive nodes, the nodes with frequency cutoffs ≤ 1, 2, 3 or 4 are removed from the selected neighborhoods.

### 6. SCZ Network validation

We compared the genes in SCZ network with dysregulated genes^84–87^ or variants^88^ identified in SCZ patients collected from recent literature, because these data have not been collected into the databases of SCZ candidate genes. The brain-specific genes were from Illumina Body Map^89^.

6.1. A total number of 2,902 brain differentially expressed genes (DEGs) (Supplementary Data 5) between schizophrenia patients and normal subjects were collected from the following literature:

1. Fromer, M. *et al.* Gene expression elucidates functional impact of polygenic risk for schizophrenia. *Nature neuroscience* **19**, 1442-1453 (2016).
2. Hoffman, G. E. *et al.* Transcriptional signatures of schizophrenia in hiPSC-derived NPCs and neurons are concordant with post-mortem adult brains. *Nature communications* **8**, 2225 (2017).
3. Sanders, A. R. *et al.* Transcriptome sequencing study implicates immune-related genes differentially expressed in schizophrenia: new data and a meta-analysis. *Translational psychiatry* **7**, e1093 (2017).
4. Chang, X. *et al.* RNA-seq analysis of amygdala tissue reveals characteristic expression profiles in schizophrenia. *Translational psychiatry* **7**, e1203, doi:10.1038/tp.2017.154 (2017).
6.2. A total number of 4,052 exome variants (Supplementary Data 5) identified in SCZ patients were collected from the following literature:

Genovese, G. *et al.* Increased burden of ultra-rare protein-altering variants among 4,877 individuals with schizophrenia. *Nature neuroscience* **19**, 1433-1441 (2016).
6.3. A total number of 2,107 brain-specific genes were collected from the following source:

*RNA-Seq of human individual tissues and mixture of 16 tissues (Illumina Body Map)*, <https://www.ebi.ac.uk/gxa/experiments/E-MTAB-513/Results>

### 7. The spatiotemporal expression of genes in human brain

Spatiotemporal co-expression was calculated as described previously (Lin et al)^90^. The brain transcriptome data was generated across 13 dissection stages (Kang et al., 2011)^91^. Since well-defined anatomical brain structures are limited during early embryonic development, the first period (4–8 postconceptional weeks, PCW) was removed from further analyses. After merging dissection stages, we defined ten partially-overlapping developmental periods ranging from 8 PCW to 40 years of age (Fig. 3a). Brain regions were grouped into four clusters using hierarchical clustering based on brain transcriptional similarity to reflect actual topological proximity and functional segregation as described in Willsey et al.^92^ As a result, 38 spatiotemporal regions were defined after eliminating one region from P2-P4 (P2-4R4) due to lack of transcriptome data. Co-expression networks were used to identify phase-specific functional relationships between genes. A co-expression between two genes was defined as positive if the pair-wise Pearson Correlation Coefficient (PCC) value was > 0.8. Using this approach, 38 different spatiotemporal co-expression networks were generated and the co-expressed gene pairs were used for the enrichment analyses of SCZ network (Fig. 3).

Development periods
| Period | Description | Intervals |
| --- | --- | --- |
| 1 | Embryonic development | $4 \leq \text{age} < 8 \text{ PCW}$ |
| 2 | Early fetal development | $8 \leq \text{age} < 10 \text{ PCW}$ |
| 3 | Early fetal development | $10 \leq \text{age} < 13 \text{ PCW}$ |
| 4 | Early mid-fetal development | $13 \leq \text{age} < 16 \text{ PCW}$ |
| 5 | Early mid-fetal development | $16 \leq \text{age} < 19 \text{ PCW}$ |
| 6 | Late mid-fetal development | $19 \leq \text{age} < 24 \text{ PCW}$ |
| 7 | Late fetal development | $24 \leq \text{age} < 38 \text{ PCW}$ |
| 8 | Neonatal and early infancy | $\text{birth} \leq \text{age} < 6\text{M}$ |
| 9 | Late infancy | $6\text{M} \leq \text{age} < 12\text{M}$ |
| 10 | Early childhood | $1 \text{ Y} \leq \text{age} < 6 \text{ Y}$ |
| 11 | Middle and late childhood | $6 \text{ Y} \leq \text{age} < 12 \text{ Y}$ |
| 12 | Adolescence | $12 \text{ Y} \leq \text{age} < 20 \text{ Y}$ |
| 13 | Young adulthood | $20 \text{ Y} \leq \text{age} < 40 \text{ Y}$ |
| 14 | Middle adulthood | $40 \text{ Y} \leq \text{age} < 60 \text{ Y}$ |
| 15 | Late adulthood | $60+ \text{ Y}$ |

### 8. Intracerebroventricular injection of serum from immune-activated mice

Intraperitoneal injection (IPI) of lipopolysaccharide (LPS) at a dosage of 150 ug/kg was done twice on post-natal day 45 and 46 to activate the immune system. The serum was taken from the immune activated mice on post-natal day 47. The intracerebroventricular injection (ICVI) of 15 ul of serum was performed on adolescent mice using microinjection. Briefly, the mice were anesthetized using 0.01ml/g of 0.75% pentobarbital sodium solution. After the hair on top of the head was trimmed, the mouse was mounted onto a stereotaxic instrument (RWD Life Science, USA). The syringe of 33 gauge (Hamilton, USA) for ICVI was washed with water for 5 times, with 70% ethanol for 5 times and then saline for 3 times. The needle was stereotaxically moved into the right lateral cerebral ventricle at anteroposterior (AP) −0.53 mm, mediolateral (ML) 0.8-0.9 mm and dorsal ventral (DV) −2.5 mm, and 15 ul of serum was injected into the lateral cerebral ventricle. All procedures were approved by the Southern Medical University Experimental Animal Ethics Committee.

### 9. Single-cell transcriptome sequencing and data analyses

#### 9.1. Single-cell RNA sequencing of the brain

The mice were anesthetized using 0.01 ml/g of 0.75% pentobarbital sodium solution and perfusion was carried out to remove blood from the brain tissues. The whole brain was taken immediately after the mouse was executed. Cell suspension of the brain was generated using MACS Adult Brain Dissociation Kit (Miltenyi Biotec, USA) following the manufacturer’s instruction. Cells were then processed through the Chromium Single-cell 3’ v2 Library Kit (10× Genomics) by the Oebiotech (Shanghai, China). Briefly, 10,000 cells were loaded onto a single channel of the 10× Chromium Controller. Messenger RNA from approximately ~7,000 cells, captured and lysed within nanoliter-sized gel beads in emulsion, was reverse transcribed and barcoded using polyA primers with unique molecular identifier sequences before being pooled, amplified, and used for library preparation. The library was then sequenced in the Illumina Nova platform.

#### 9.2. Identifying different types of cell population

Demultiplexing of the bcl file into a FASTQ file was performed using bcl2fastq (version 2.19.0). We generated the UMI count matrix with cellranger’s^93^ (version 2.2.0) ‘cellranger count’ function for each library, using mouse (mm10-1.2.0) genome reference sequences. Dimensionality reduction of data was performed by principal component analysis (PCA) (N = 10 principal components) using R toolkit Seurat’s (version 2.3.4) RunTSNE function^94^, and the reduced data were visualized in two dimensions using the *t*-SNE non-linear dimensionality reduction method. Clustering for expression similarity was performed using a shared nearest neighbor (SNN) modularity-optimization-based clustering algorithm (with dimension = 10) by Seurat’s FindClusters function. We use the specific markers of different types of cells in the mouse brain for identification of the cell population^95^.

#### 9.3. Monocle differential expression analysis

Cells belonging to the astrocyte, microglia, neuron, oligodendrocyte as 4 groups depicted in Fig. 5d were used for DE analysis. Gene-cell matrices produced by Seurat were loaded into R and DE analysis was performed with monocle (version 2.10.1)^96^. DEGs were found using Monocle’s DifferentialGeneTest function and the genes with *q*-value < 0.01 are kept.

