## Supplemental figure for "Systematic search for schizophrenia pathways sensitive to perturbation by immune activation"

a

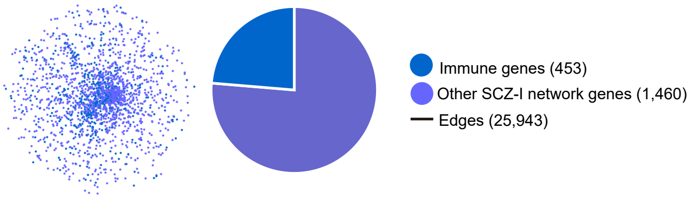


SCZ-i subnetwork at early fetal developmental stages (P02-04)

b

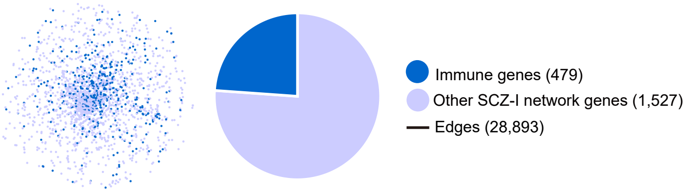


SCZ-i subnetwork at mid-late fetal developmental stages (P05-06)

c

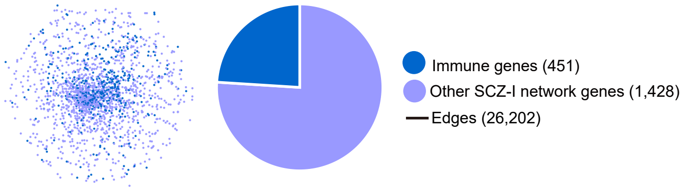


SCZ-i subnetwork at adolescent developmental stages (P12-13)

**Supplementary Fig. 1. The SCZ-i subnetworks in three different developmental stages (Corresponding to Fig. 4).**

a. Construction of the SCZ-i subnetwork in early fetal developmental stage. Co-expressed genes in early fetal developmental stage (P02-04) were mapped onto the SCZ-i network and the SCZ-i subnetwork in this stage was extracted.

b. Construction of the SCZ-i subnetwork in mid-late fetal developmental stage. Co-expressed genes in mid-late developmental stage (P05-06) were mapped onto the SCZ-i network and the SCZ-i subnetwork in this stage was extracted.

c. Construction of the SCZ-i subnetwork in adolescent stage. Co-expressed genes in adolescent stage (P12-13) were mapped onto the SCZ-i network and the SCZ-i subnetwork this stage was extracted.

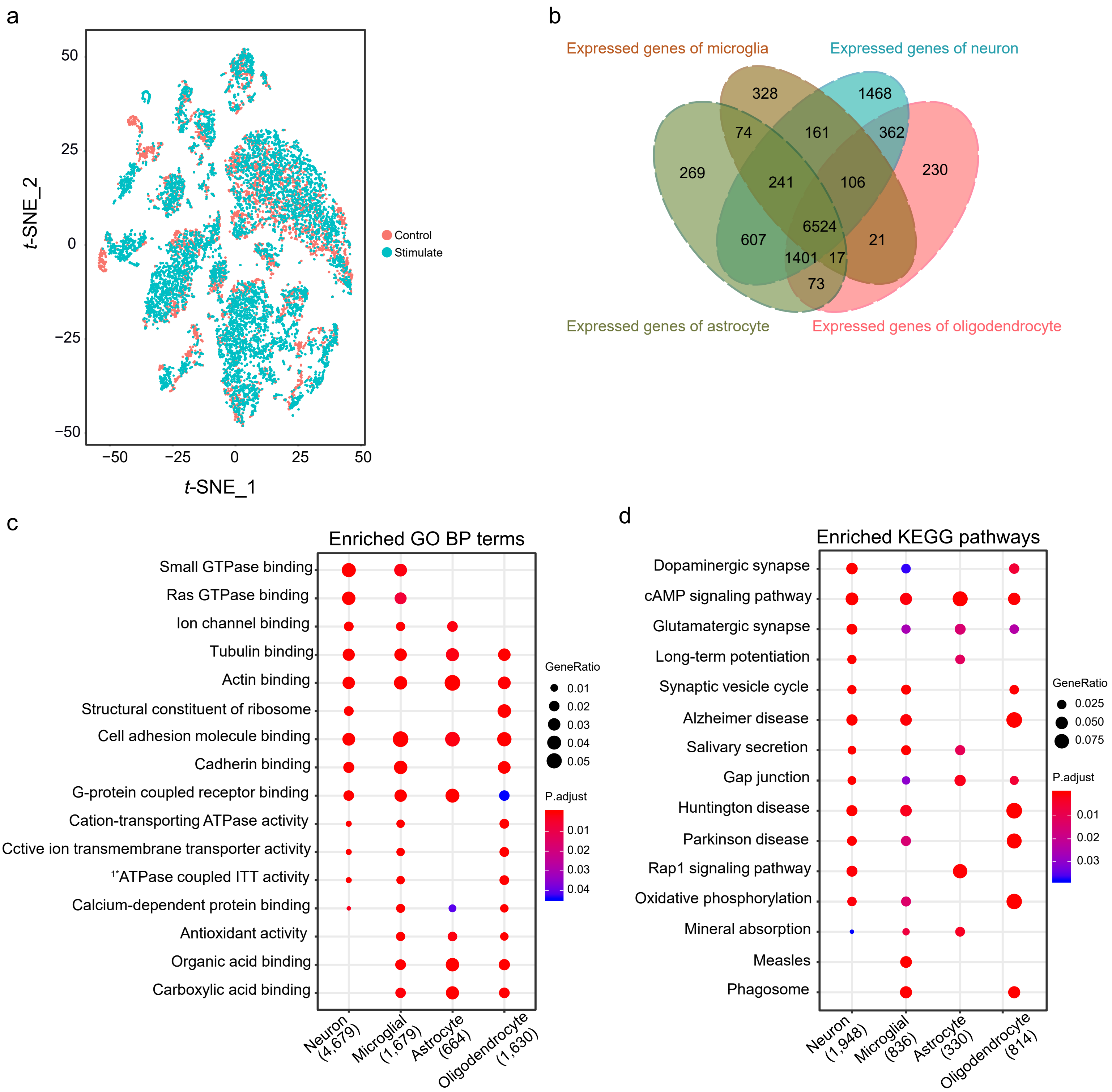


**Supplementary Fig. 2. Singe-cell transcriptome profiling on immune-activated brains of adolescent mice reveals key pathways linking immune activation to synapse remodeling (Corresponding to Fig. 5).**

a. The t-SNE plot constructed from the normalized and corrected logarithm of expression values of highly variable genes in cells. Red represents brain cells from mice without immune activation; cyan represents brain cells from mice with immune activation.

b. Venn diagram for expressed genes in 4 major cell types in the brain.

c-d. The GO terms and KEGG pathways with overrepresented DEGs of neurons, microglia, astrocytes and oligodendrocytes. Top 15 highly enriched GO terms (c) or pathways (d) are shown as dots. The sizes of the dots represent ratios of DEGs in pathways and the colors represent the adjusted p-values. The numbers under each stage are the numbers of identified genes in each category. See also Supplementary Data 8 for the complete list of enriched GO terms and KEGG pathways.

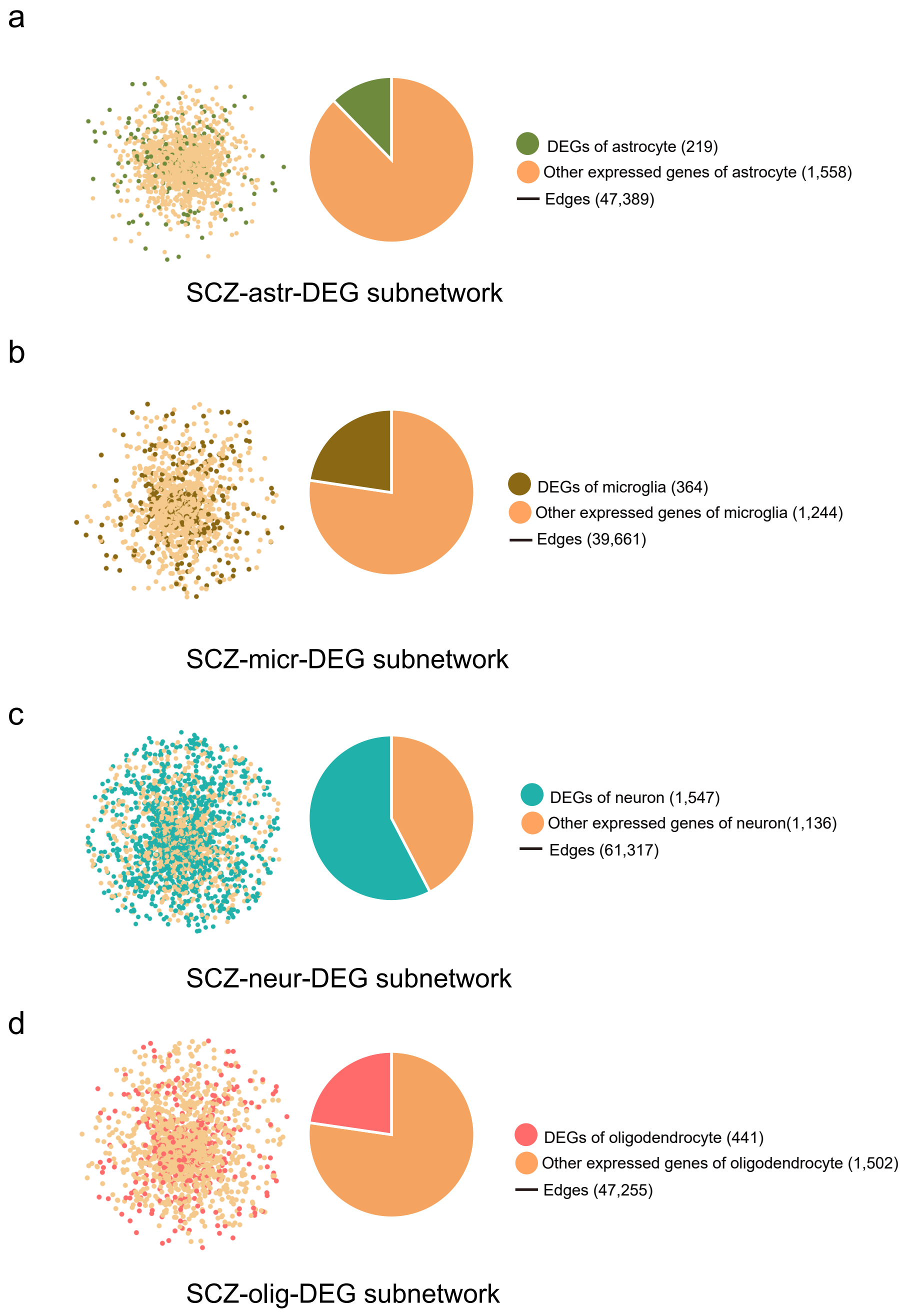


**Supplementary Fig. 3. SCZ-DEG subnetworks of different cell types (Corresponding to Fig. 5).**
The expressed genes of each cell-type were mapped onto the SCZ Network, and the SCZ-cell-type subnetworks were extracted; and then DEGs of each cell type in the immune-activated brain were mapped onto the SCZ-cell-type subnetworks and the SCZ-cell-type-DEG subnetworks were extracted to include DEGs and their first-degree neighbors.

a. Construction of SCZ-astrocyte-DEG subnetwork (SCZ-astr-DEG subnetwork). Astrocyte expressed genes were mapped onto the SCZ Network and SCZ-astrocyte subnetwork was extracted; then the DEGs of astrocytes in the immune-activated brain were mapped onto SCZ-astrocyte subnetwork and the SCZ-astr-DEG subnetwork was extracted to include the DEGs of astrocytes and their first-degree neighbors.
b. Construction of SCZ-microglia-DEG subnetwork (SCZ-micr-DEG subnetwork). Microglia expressed genes were mapped onto the SCZ Network and SCZ-microglia subnetwork was extracted; then the DEGs of microglia in the immune-activated brain were mapped onto SCZ-microglia subnetwork and the SCZ-micr-DEG subnetwork was extracted to include the DEGs of microglia and their first-degree neighbors.
c. Construction of SCZ-neuron-DEG subnetwork (SCZ-neur-DEG subnetwork). Neuron expressed genes were mapped onto the SCZ Network and SCZ-neuron subnetwork was extracted; then the DEGs of neurons in the immune-activated brain were mapped onto SCZ-neuron subnetwork and the SCZ-neur-DEG subnetwork was extracted to include the DEGs of neurons and their first-degree neighbors.
d. Construction of SCZ-oligodendrocyte-DEG subnetwork (SCZ-olig-DEG subnetwork). Oligodendrocyte expressed genes were mapped onto the SCZ Network and SCZ-oligodendrocyte subnetwork was extracted; then the DEGs of oligodendrocytes in the immune-activated brain were mapped onto SCZ-oligodendrocyte subnetwork and the SCZ-olig-DEG subnetwork was extracted to include the DEGs of oligodendrocytes and their first-degree neighbors.
