## Supplemental information for "Systematic search for schizophrenia pathways sensitive to perturbation by immune activation"

### CONTENTS

#### Supplementary Figures

Supplementary Fig. 1. The SCZ-i subnetworks in three different developmental stages (Corresponding to Fig. 4).

Supplementary Fig. 2. Singe-cell transcriptome profiling on immune-activated brains of adolescent mice reveals key pathways linking immune activation to synapse remodeling (Corresponding to Fig. 5).

Supplementary Fig. 3. SCZ-cell-type-DEG subnetworks (Corresponding to Fig. 5).

#### Supplementary Data (Excel tables)

Supplementary Data 1. SCZ-candidate genes and immune genes collected from databases and literature.

Supplementary Data 2. Risk-score or Guilt-score assigned to genes in a comprehensive high quality human interactome.

Supplementary Data 3. SCZ-related neighborhoods at different cutoffs.

Supplementary Data 4. The final SCZ network.

Supplementary Data 5. Genetic variants and differentially expressed genes collected from literature for validation of the SCZ Network.

Supplementary Data 6. Expression data from 4 regions of the brain and Overrepresentation of SCZ Network genes in spatiotemporal phases.

Supplementary Data 7. The SCZ-i Network

Supplementary Data 8. Differentially expressed genes identified by single cell sequencing

Supplementary Data 9. SCZ-cell-type-DEG subnetworks and functional enrichment.

### Supplementary Data

**(Excel files)**

#### Supplementary Data 1. SCZ-candidate genes and immune genes collected from databases and literature.

Sheet 1. SCZ-candidate genes

Sheet 2. Immune genes

#### Supplementary Data 2. Risk-score or Guilt-score assigned to the genes in a comprehensive high-quality human interactome.

Sheet 1. Human interactome data (nodes)

Sheet 2. Human interactome data (edges)

Sheet 3. R-scores and G-scores

Sheet 4. Nodes distance and edge weight

#### Supplementary Data 3. SCZ-related network neighborhoods at different cutoffs.

Sheet 1. SCZ-related network neighborhoods at seed ratio >85%

Sheet 2. SCZ-related network neighborhoods at seed ratio >90%

#### Supplementary Data 4. The SCZ Network.

Sheet 1. The SCZ Network (nodes)

Sheet 2. The SCZ Network (edges)

#### Supplementary Data 5. Genetic variants and differentially expressed genes of schizophrenic brains collected from literature for validation of the SCZ Network.

Sheet 1. Brain differentially expressed genes (DEGs) in schizophrenic brains

Sheet 2. Exome variants identified in SCZ patients

Sheet 3. Enrichment results

Sheet 4. Tissue-specific genes

#### Supplementary Data 6. Expression data from 4 regions of the brain and overrepresentation of SCZ Network genes in spatiotemporal phases.

Sheet 1. Co-expressed genes from 4 regions of the brain at different periods

Sheet 2. Overrepresentation of the SCZ Network genes in spatiotemporal phases

Sheet 3. Overrepresentation of non-immune genes of the SCZ Network in spatiotemporal phases

Sheet 4. Overrepresentation of immune genes of the SCZ Network in spatiotemporal phases

#### Supplementary Data 7. The SCZ-i network

Sheet 1. The SCZ-i subnetwork (nodes)

Sheet 2. The SCZ-i subnetwork (edges)

Sheet 3. GO terms enriched with genes in the SCZ-i Subnetwork

Sheet 4. KEGG pathways enriched with genes in the SCZ-i Subnetwork

Sheet 5. GO terms enriched with genes in the SCZ-i Subnetwork at different brain developmental stages

Sheet 6. KEGG pathways enriched with genes in the SCZ-i Subnetwork at different brain developmental stages

#### **Supplementary** Data 8. Differentially expressed genes in brain cells identified by single-cell sequencing

Sheet 1. Overrepresentation of genes of the SCZ Network in 9 different cell populations

Sheet 2. All expressed genes

Sheet 3. Differentially expressed genes of astrocytes

Sheet 4. Differentially expressed genes of microglia

Sheet 5. Differentially expressed genes of neurons

Sheet 6. Differentially expressed genes of oligodendrocytes

Sheet 7. GO terms enriched with DEGs of 4 cell populations

Sheet 8. KEGG pathways enriched with DEGs of 4 cell populations

#### Supplementary Data 9. SCZ-cell-type-DEG subnetworks and functional enrichment.

Sheet 1. The SCZ-astrocyte-DEG subnetwork (nodes)

Sheet 2. The SCZ-astrocyte-DEG subnetwork (edges)

Sheet 3. The SCZ-microglia-DEG subnetwork (nodes)

Sheet 4. The SCZ-microglia-DEG subnetwork (edges)

Sheet 5. The SCZ-neuron-DEG subnetwork (nodes)

Sheet 6. The SCZ-neuron-DEG subnetwork (edges)

Sheet 7. The SCZ-oligodendrocyte-DEG subnetwork (nodes)

Sheet 8. The SCZ-oligodendrocyte-DEG subnetwork (edges)

Sheet 9. GO terms enriched with genes in SCZ-cell-type-DEG subnetworks

Sheet 10. KEGG pathways enriched with genes in SCZ-cell-type-DEG subnetworks
